# Noninvasive prenatal test of methylmalonic academia cblC type through targeted sequencing of cell-free DNA in maternal plasma

**DOI:** 10.1101/425918

**Authors:** Lianshu Han, Chao Chen, Fengyu Guo, Jun Ye, Zhiyu Peng, Wenjuan Qiu, Yaoshen Wang, Wei Li, Huiwen Zhang, Lili Liang, Yu Wang, Huanhuan Wang, Xing Ji, Jun Sun, Xuefan Gu

**Author notes:** The first two authors contributed equally to this work. **Corresponding author:** Xuefan Gu and Jun Sun: Xinhua Hospital, Shanghai Institute for Pediatric Research, Shanghai Jiao Tong University School of Medicine. 1665Kongjiang Road, Shanghai 200092, China. : Tianjin Medical Laboratory, E3 Building, Airport Economics Zone, Tianjin 300308, China.

## Abstract

Methylmalonic acidemia (MMA) cblC type is the most frequent inborn error of intracellular cobalamin metabolism which is caused by mutations of *MMACHC* gene. Non-invasive test of MMA for pregnant women facilitates safe and timely prenatal diagnosis of the disease. In our study, we aimed to design and validate a haplotype-based noninvasive prenatal test (NIPT) method for cblC type of MMA. Targeted capture sequencing using customized hybridization was performed utilizing gDNA (genomic DNA) of trios including parents and an affected proband to determine parental haplotypes associated with the mutant and wild allele. The fetal haplotype was inferred later based on the high depth sequencing data of maternal plasma as well as haplotype linkage analysis. The fetal genotypes deduced by NIPT were further validated by amniocentesis. Haplotype-based NIPT was successfully performed in 21 families. The results of NIPT of 21 families were all consistent with invasive prenatal diagnosis, which was interpreted in a blinded fashion. Three fetuses were identified as compound heterozygosity of *MMACHC*, 9 fetuses were carriers of *MMACHC* variant, and 9 fetuses were normal. These results indicated that the haplotype-based NIPT for MMA through small target capture region sequencing is technically accurate and feasible.

## Introduction

Methylmalonic acidemia (MMA) is caused by a deficiency of methylmalonyl-coA mutase or coenzyme adenosylcobalamin (AdoCbl). The cblC type combined with methylmalonic acidemia and homocystinuria is a kind of autosomal recessive hereditary disease and the incidence was found to be in 1 in 48,000 to 1 in 250,000 worldwide (Carrillo-Carrasco *et al*. 2012; Wang *et al*. 2010). Newborn screening for MMA in Shandong province of China showed an estimated prevalence of 1 in 3920 during birth (Han et al. 2016). The MMA cblC type caused by mutations in the *MMACHC* gene (NM_015506.2) located on chromosome 1p34.1 is the most frequent congenital error in intracellular co-amine metabolism. Prenatal diagnosis of MMA was essential due to age of onset in neonatal period and serious symptoms such as multiple system damage and lethal possibility. It can not only contribute to early medical management of infants, but also allows treatment of affected foetus as soon as possible to reduce irreversible organ damage associated with metabolic acidosis and high blood ammonia. (Huemer *et al*. 2005; Trefz *et al*. 2016).

Recently, several noninvasive prenatal test (NIPT) methods aimed to avoid miscarriage or infection risk method have been developed (Evans *et al*. 2002; Mujezinovic and Alfirevic 2007). Besides large-scale clinical application in fetal aneuploidies screening, NIPT for dominant single-gene disorders confined to detect paternally inherited and de novo mutations has also been introduced to clinical trials (Lo *et al*. 1998; Fan *et al*. 2008). However, NIPT for recessive single gene disorders are still at laboratory research stage. Our study group has developed a haplotype-based method of NIPT for recessive inherited single gene diseases using linkage analysis of trios’ members and has been validated in several single gene disorders (Xu *et al*. 2015; Meng *et al*. 2014; Lam *et al*. 2012; Ma *et al*. 2014; Ye *et al*. 2018; Chen *et al*. 2017). The feasibility and accuracy of this method depends on the informative SNPs used to construct the haplotype and the selection of the targeted region. NIPT of MMA based on haplotype analysis method has not been reported yet, and so this study has been designed.

In our study, the NIPT of MMA can be accomplished using a 141.39 kb customized probe, including *MMACHC* and 1125 surrounding highly heterozygous SNPs (0.3<minor allele frequency<0.5) distributed within the 1 Mb on chromosome 1 (Figure 1). Our study indicated that the haplotype-based NIPT for MMA through small target capture region sequencing is technically accurate and feasible, and also further highlighted the feasibility of NIPT of monogenic diseases.

**Figure 1.**
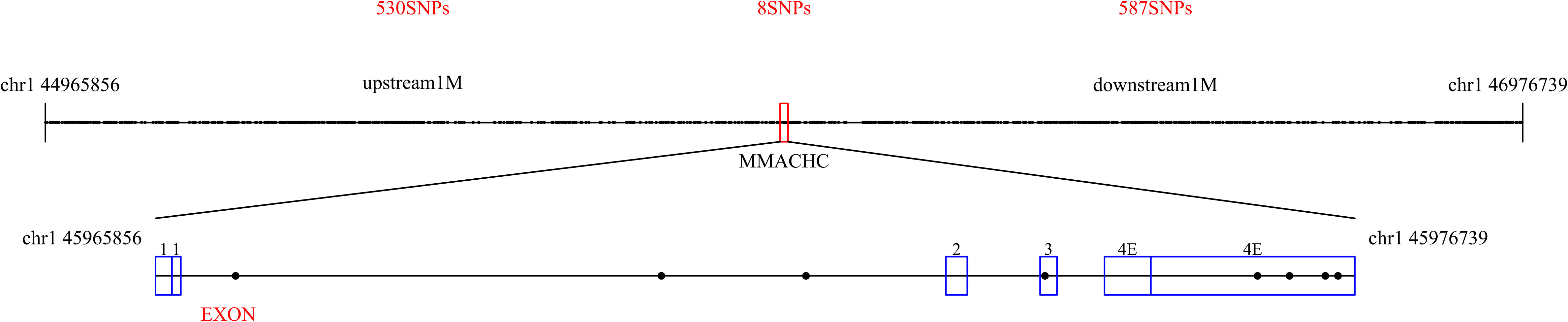
Target region of *MMACHC* gene and SNPs used for haplotyping. Custom NimbleGen probes within the 141.39 kb region was designed including the *MMACHC* gene coding region and 1125 surrounding highly heterozygous SNPs distributed within 1 Mb region.

## Materials and Methods

### Patient recruitment

Twenty-one at-risk families including singleton pregnant woman, her husband, and proband diagnosed with MMA were enrolled at Shanghai Xin Hua Hospital with genetic counseling and informed consent. The institutional review board (IRB) of BGI and Shanghai Xinhua Hospital approved our study with approval number BGI-IRB NO. FT15195 and XHEC-C-2015-025.

Ten causative mutations in four exons of *MMACHC* gene in probands and their parents were already identified prior to performing NIPT of MMA during pregnancy. Before amniocentesis, blood samples were drawn from each pregnant woman at 16–20 weeks of gestation and her family members was used for NIPT of MMA. The maternal plasma should be separated as soon as possible. Amniotic fluid (AF) samples for routine prenatal diagnosis were obtained at 16–18 weeks of gestation. The data analysis of fetal DNA sample was blinded to NIPT. The AF samples of F01-F06 family were also sent to BGI to evaluate the accuracy of inferred paternal and maternal specific loci of fetus compared with the AF standard haplotype.

### Targeted sequencing

The extraction of gDNA and maternal plasma DNA was accomplished using the commercial kits from QIAGEN. The NGS library of gDNA was constructed according to the Illumina standard protocol. The construction of cell-free DNA library as a result of its micro inputs nature was performed by the Kapa Biosysterm library preparation kit. The hybrid captures of gDNA and cf-DNA libraries were separately carried out using the same probe. The post-capture libraries were sequenced using PE 101 bp on Illumina platform (Hiseq 2500).

### Variation calling

The raw data were aligned to the human reference sequence (Hg19, GRCh37) by BWA software (0.7.12) in the paired end mode. After removal of low-quality reads including duplicated reads and multiple aligned reads using Picard Tools, Variation calling was accomplished through GATK software. Only the variations with depth greater than 50x will be analyzed in the next step.

### Estimation of fetal concentration and plasma sequencing error

The fetal genotype should be heterozygous state with different homozygous genotype of parents according to the Mendel’s laws. So, it could be estimated as two times of the percentage of the minor allele depth to the total depth of this allele. The sequencing error could be described as the ratio of the count of different loci reads to total count of this SNP reads when the genotypes of parents are same homozygous.

### NIPT for MMA

Haplotypes linked with wild and mute allele were constructed using SNP information within flanking and coding region of *MMACHC* gene (You et al. 2014). Hap 0 was defined as the pathogenic haplotype and Hap1 was defined as the wild-type haplotype. The fetal inheritance from father was determined using the set of SNPs that were heterozygous in father but homozygous in mother. SNPs that were heterozygous in mother but homozygous in father were used to determine fetal inheritance from mother. Hidden Markov Model (HMM) and Viterbi algorithm was used to deduce the fetal haplotypes using the target region data of plasma (Ma *et al*. 2017).

### Accuracy of NIPT for MMA

The Sanger sequencing and standard haplotypes of fetal gDNA obtained using amniotic fluid was further operated to validate the uniformity of NIPT result.

## Results

### Sequencing data of recruited families

Sanger sequencing of *MMACHC* was operated to determine the variation of pathogenic mutation (Table 1). The mean depth of gDNA and cf-DNA was about 147.48x (63.41x–348.05x) and 237.97x (89.67x396.23x), respectively. The coverage with more than 20x was approximately 98.14% (90.09–99.68%). The mean capture efficiency and duplicate rate was 50.19% (27.12%–75.23%) and 7.37% (4.98%–44.09%) in all the samples (Table S1).

**Table 1.**
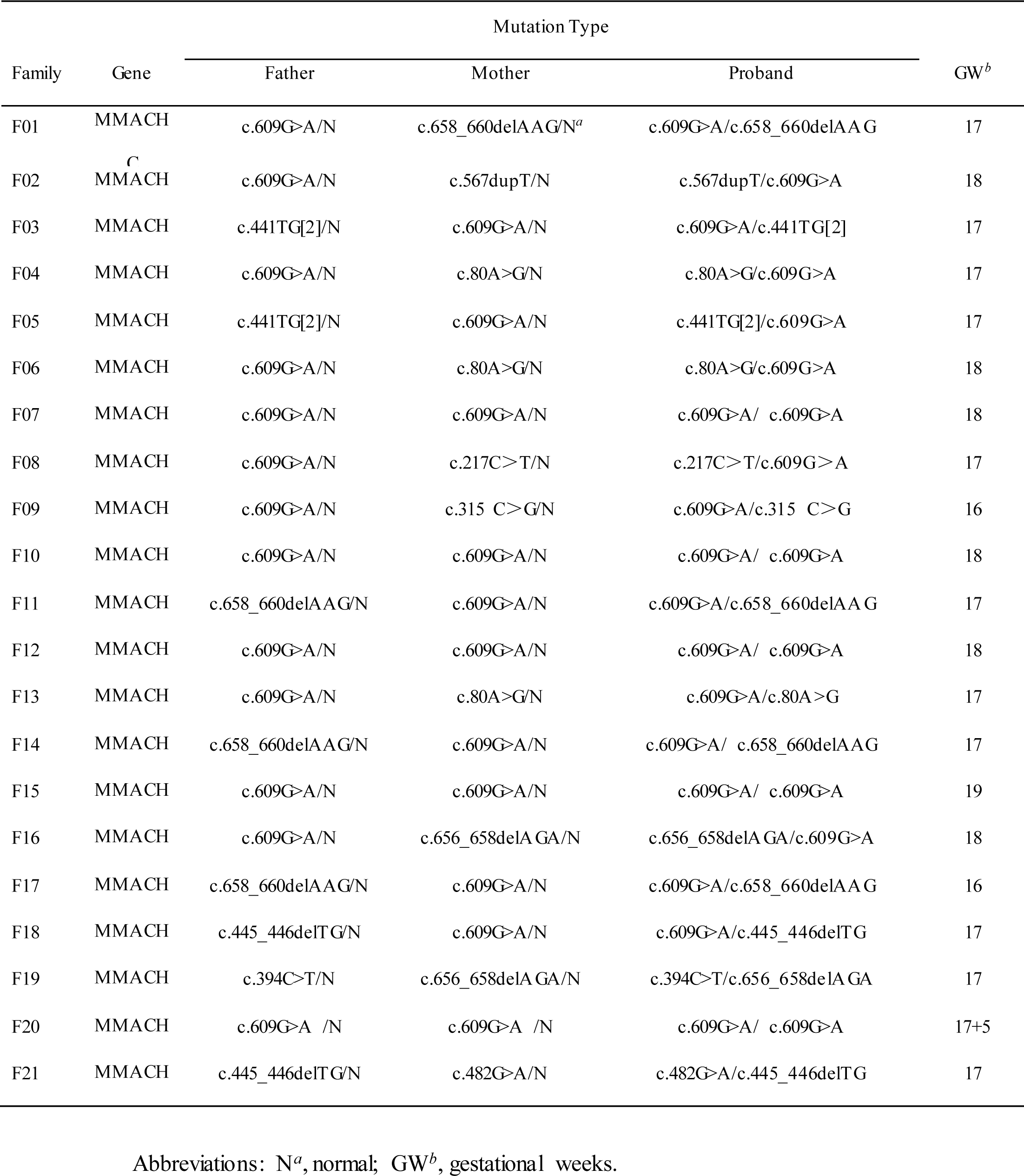
Molecular Diagnosis Results in 21 Families.

### Estimation of fetal concentration and plasma sequencing error

For these twenty-one pregnant women, the cff-DNA concentrations varied from 4.62% to 18.96% during the second trimester (Table 2), showing significant differences between the individuals. The mean sequencing error rate of plasma was 0.41% (0.02–1.18%) (Table 2). The data suggested high experimental quality for next analysis.

**Table 2.** Statistics of Error Rate and Fetal DNA Fraction.

| **Pedigree** | **Type 1 SNP<sup>a</sup>** | **Type 2 SNP<sup>b</sup>** | **Error Rate** | **Fetal DNA fraction** |
| --- | --- | --- | --- | --- |
| O1 | 86 | 5 | 0.0200% | 7.64% |
| F02 | 104 | 463 | 1.1788% | 11.45% |
| F03 | 79 | 430 | 0.1936% | 16.23% |
| F04 | 107 | 215 | 0.2437% | 9.07% |
| F05 | 85 | 14 | 0.3360% | 7.66% |
| F06 | 99 | 43 | 1.0153% | 18.96% |
| F07 | 125 | 1 | 0.1631% | 14.89% |
| F08 | 66 | 421 | 0.7156% | 10.93% |
| F09 | 80 | 23 | 0.0941% | 4.62% |
| F10 | 110 | 61 | 0.4655% | 11.41% |
| F11 | 115 | 4 | 0.2434% | 17.84% |
| F12 | 82 | 149 | 0.3810% | 6.85% |
| F13 | 329 | 39 | 0.2170% | 11.70% |
| F14 | 64 | 24 | 0.4847% | 9.86% |
| F15 | 66 | 7 | 0.5687% | 11.66% |
| F16 | 70 | 165 | 0.3680% | 10.77% |
| F17 | 71 | 441 | 0.2319% | 13.03% |
| F18 | 49 | 166 | 0.8472% | 10.52% |
| F19 | 238 | 141 | 0.2406% | 5.90% |
| F20 | 66 | 23 | 0.3877% | 5.85% |
| F21 | 143 | 8 | 0.2170% | 8.20% |

SNPs<sup>a</sup> that were homozygous with the same type of parents but different bases in the plasma, which were used to calculate the sequencing error rate of plasma; SNPs<sup>b</sup> that were homozygous in both parents but different types, which were used to calculate fetal DNA fractions.

### Construction of Parental haplotype

The parental haplotype was constructed by the genotyping information from a trio strategy of father, mother and proband. Clean data with more than 20-fold coverage were about97.88% and about 1041 SNPs on the target region were detected.

### Noninvasive prenatal diagnosis of fetal MMA

The number of SNPs identified ranged from 854 to 1211. The number of informative SNPs which were used to predict the combination of fetal haplotype inherited from mother and father were 122 (20–323) and 113 (24–290). Parental haplotypes were successfully constructed in F01 family. 155 SNPs was used to determine that the fetus inherited wild allele from the father and none SNP supported that fetus inherited the pathogenic haplotype. 86 SNPs was used to determine that the fetus inherited wild allele from the mother and none SNP supported that fetus inherited the pathogenic haplotype. So, the fetus of F01 was normal because of the F1+M1 haplotype (Table 3 and Figure 2). Based on this strategy, nine fetuses were diagnosed as carriers, nine fetuses were normal and three fetuses were affected by cblC type of MMA due to the compound heterozygous mutation of *MMACHC* (Table 3 and Figure 2).

**Table 3.** The NIPT Results of 21 Studied Families.

| Family | Gene | SNPs For F0<sup>a</sup> | SNPs For F1<sup>a</sup> | SNPs For M0<sup>a</sup> | SNPs For M1<sup>a</sup> | Fetal Haplotype | Fetal Genotype | NIPT Results | Invasive Test Results | Consistency<sup>b</sup> |
| --- | --- | --- | --- | --- | --- | --- | --- | --- | --- | --- |
| F01 | MMACHC | 0 | 155 | 0 | 86 | F1+M1 | N | Normal | N | Y<sup>c</sup> |
| F02 | MMACHC | 37 | 0 | 0 | 86 | F0+M1 | c.609G>A/N | Carrier | c.609G>A/N | Y |
| F03 | MMACHC | 0 | 60 | 73 | 0 | F1+M0 | c.609G>A/N | Carrier | c.609G>A/N | Y |
| F04 | MMACHC | 0 | 42 | 94 | 0 | F1+M0 | c.80A>G/N | Carrier | c.80A>G/N | Y |
| F05 | MMACHC | 109 | 0 | 122 | 0 | F0+M0 | c.441TG[2]/c.609G>A | affected | c.441TG[2]/c.609G>A | Y |
| F06 | MMACHC | 0 | 325 | 20 | 0 | F1+M0 | c.80A>G/N | Carrier | c.80A>G/N | Y |
| F07 | MMACHC | 0 | 140 | 0 | 91 | F1+M1 | N | Normal | N | Y |
| F08 | MMACHC | 129 | 0 | 42 | 0 | F0+M0 | c.217C>T/c.609G>A | affected | c.217C>T/c.609G>A | Y |
| F09 | MMACHC | 0 | 160 | 77 | 0 | F1+M0 | c.315 C>G/N | Carrier | c.315 C>G/N | Y |
| F10 | MMACHC | 0 | 24 | 0 | 273 | F1+M1 | N | Normal | N | Y |
| F11 | MMACHC | 0 | 81 | 0 | 149 | F1+M1 | N | Normal | N | Y |
| F12 | MMACHC | 0 | 43 | 203 | 0 | F1+M0 | c.609G>A/N | Carrier | c.609G>A/N | Y |
| F13 | MMACHC | 0 | 26 | 0 | 117 | F1+M1 | N | Normal | N | Y |
| F14 | MMACHC | 220 | 0 | 96 | 0 | F0+M0 | c.609G>A/c.658-660delAAG | affected | c.609G>A/c.658-660delAAG | Y |
| F15 | MMACHC | 0 | 27 | 86 | 0 | F1+M0 | c.609G>A/N | Carrier | c.609G>A/N | Y |
| F16 | MMACHC | 290 | 0 | 0 | 111 | F0+M1 | c.609G>A/N | Carrier | c.609G>A/N | Y |
| F17 | MMACHC | 0 | 109 | 0 | 84 | F1+M1 | N | Normal | N | Y |
| F18 | MMACHC | 0 | 54 | 0 | 323 | F1+M1 | N | Normal | N | Y |
| F19 | MMACHC | 108 | 0 | 0 | 128 | F0+M1 | c.394C>T/N | Carrier | c.394C>T/N | Y |
| F20 | MMACHC | 0 | 27 | 0 | 230 | F1+M1 | N | Normal | N | Y |
| F21 | MMACHC | 0 | 274 | 0 | 73 | F1+M1 | N | Normal | N | Y |

<sup>a</sup>SNPs for F/M represent the number of SNP supporting the inheritance of fetal haplotype from parents. F0/M0, fetal-inherited mutant haplotype; F1/M1, fetal-inherited normal haplotype; <sup>b</sup>Consistency represents the comparison of NIPT results with invasive testing results; <sup>c</sup> Y, yes; N, normal.

**Figure 2.**
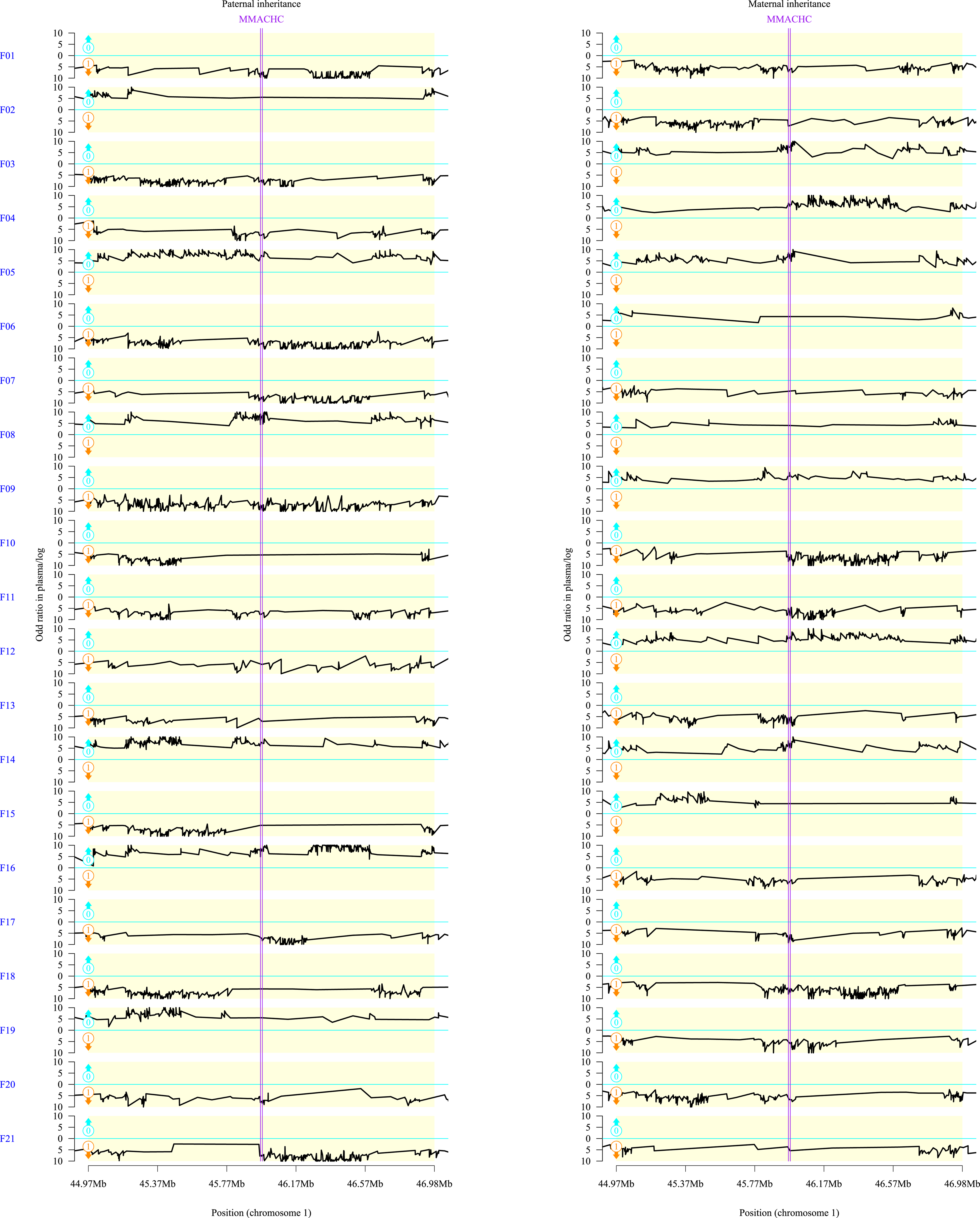
Fetal Haplotype Prediction. X-axis represents the locus on target region; Y-axis represents the logarithm of the ratios of different fetal haplotype combinations. The black lines above zero (cyan lines) indicate that the fetus inherited the pathogenic haplotype (Hap0), and the black lines below zero indicate that the fetus inherited the normal haplotype (Hap1). Two purple vertical lines represent the region of the *MMACHC* gene. Left chart and right chart represents the results of fetal-inherited paternal haplotypeand maternal haplotype, respectively.

### Accuracy of NIPT for MMA

The Sanger results of fetal DNA were 100% consistent with NIPT result (Table 2) and SNPs deduced using NIPT were 100% consistent with standard haplotypes of fetal gDNA in F01 to F06 (Table S2).

## Discussion

The cumulative incidence of monogenic disease accounted for 7% of birth defects, while the chromosomal abnormalities accounted for 6%according to the March of Dimes Global Report on Birth Defects 2006. A growing number of birth defects and diseases can be diagnosed prenatally and treated before birth in some cases. With the large-scale clinical application of NIPT for fetal aneuploidies based on massively parallel sequencing approach (Benn *et al*. 2012; Norton *et al*. 2012; Dar *et al*. 2014), NIPT of monogenic disease remains the next frontier. In the recent years, NIPT of monogenic disorders has been actively investigated and it has a great clinical application prospect, providing requisite prognostic data for clinical intervention. Initially, NIPT of monogenic disorders relied on the detection or exclusion of paternally inherited mutations or de novo mutations by direct detection method based on PCR. PCR-based method such as ddPCR (Lun *et al*. 2008), QPCR (Guissart *et al*. 2017), PCR-RED (Chitty *et al*. 2015) and cSMART (Chen *et al*. 2016; Han *et al*. 2017) have been reported in β thalassemia, cystic fibrosis, thanatophoric dysplasia, Wilson disease and autosomal recessive nonsyndromic hearing loss, respectively. PCR-based method involves the advantage of simple operation. However, the design of primer or probe at the mutation site that is confined only to SNP and indel remains difficult. NIPT based on PCR cannot be applied to the major mutation type with copy number variance (CNV). The sensitivity and specificity of PCR-based NIPT is affected by fetal DNA fraction and quality of sample.

Maternally inherited alleles and alleles shared by both parents were detected by more sophisticated techniques such as haplotype-based strategy called indirect method. Haplotype-based strategy such as relative haplotype dosage (RHDO) (Lam *et al*. 2012) and proband-assisted haplotype phasing have been successfully reported in several recessive monogenic diseases. These historical data were retrospectively analyzed in our study (Table 4). Fifty-seven high risk families who had an affected child were enrolled in the clinical research to validate the accuracy and feasibility of NIPT. All the NIPT results were consistent with invasive prenatal diagnosis. Overall, the data obtained from all the 57 families included in the test have shown sensitivity and specificity rates of 100%, with 0% failure rate. The accuracy of inferred fetal maternal alleles and paternal alleles using plasma sequencing data were almost 100% compared with the standard haplotype obtained by the AF data without considering the recombination point. Other experiments were necessary to determine the precise positioning of recombination and whether the proband or fetus involve the recombination. Ye *et al*. (2018) showed that when the plasma sequence depth was 200X, the accuracy of fetal inherited maternal haplotype was above 99% and when the fetal fraction was between 5% and 10%, the mean number of SNP was about 20 to reach 99% detection accuracy. Previous data showed that the haplotype-based NIPT can be applied to genes with highly homologous sequences like CYP21A2 and SMN1, which is almost impossible to be detected by directly sequencing the pathogenic mutations.

**Table 4.** Studies Reporting on NIPT of Monogenic Disease.

| Author | Method | Disease | Case No. | NIPT Results | Consistency | Accuracy of maternal alleles | Accuracy of paternal alleles |
| --- | --- | --- | --- | --- | --- | --- | --- |
| Dennis Lo | RHDO | β-thalassemia | 2 | 2 Carriers | Y | - | - |
| Dennis Lo | RHDO | CAH | 14 | 5 Carriers+2 Normal+7 affected | Y | - | - |
| Yanqin You | PAHP | MSUD | 1 | affected | Y | - | - |
| Zhengfeng Xu | PAHP | CAH | 1 | Carriers | Y | 96.41% | 97.81% |
| Meng Meng | PAHP | GJB2 | 1 | Carriers | Y | - | - |
| Yan Xu | PAHP | DMD | 8 | 1 Carriers+2 Normal+5 affected | Y | 99.98% | - |
| Min Chen | PAHP | SMA | 5 | 2 Carriers+2 Normal+1 affected | Y | - | - |
| Zhengfeng Xu | PAHP | CAH | 12 | 6 Carriers+6 affected | Y | 100% | 100% |
| Xuefan Gu | PAHP | PKU | 13 | 4 Carriers+4 Normal+5 affected | Y | 100% | 100% |
| **Total** | - | - | 57 | 22 Carriers+10 Normal+25 affected | Y |  |  |

RHDO, relative haplotype dosage; PAHP, proband assisted haplotype phasing; consistency means whether the NIPT result was consistent with the invasive prenatal testing, Y represents yes.

MMA is one of the most common disorders of congenital organic acid metabolism. Early prenatal treatment may have an impact on the long-term complications associated with cblC disease. Prior studies have evaluated the effects of prenatal HOCbl administration and the results showed a decrease in the maternal metabolites (Huemer *et al*. 2005). Although the haplotype-based strategy was successfully implemented in several autosomal recessive disorders, this has not been applied in MMA till now. The feasibility and accuracy of this method depends on the informative SNPs used to construct the haplotype and the selection of the targeted region, and hence investigation of technical feasibility is necessary for each target disease. Our study initially demonstrated the feasibility of haplotype-based NIPT for *MMACHC*. In our study, the fetal genotype of 21 families was successfully determined using haplotype-assisted NIPT. The result of this study was consistent with the findings of invasive method. Based on these data, the feasibility and accuracy of NIPT of MMA using haplotype strategy has been demonstrated. The sensitivity and specificity rates were both 100%, with 0% failure rate. The accuracy of the fetal alleles inherited from parents deduced by haplotype strategy were almost 100% compared with the standard haplotype obtained by the AF data in families from F01 to F06 (Table S2). The other family results showed that the haplotype-based NIPT method allowed 100% concordance with the invasive diagnostic approach.

In most of the reported cases, the targeted region is always relatively large, and the sequencing cost remained exorbitant, hampering the clinical utility of this technology. In this research, we have demonstrated the feasibility of NIPT of MMA by sequencing a much smaller region. This method requires trio members of the family for constructing the parental haplotype linked to the mutation allele. The sensitivity and specificity of this method have been reported to be 100% validated by the invasive result, which is the greatest advantage of haplotype-based strategy NIPT. A fatal flaw of the haplotype-based strategy is that the feasibility is highly depend on the availability of proband sample.

Linked reads method like 10xGenomics to specify the individual haplotype directly without proband has been recently reported by Dennis Lo in NIPT of monogenic diseases (Hui *et al*. 2017), but the technology has several limitations. The most important restrictive factor for clinical application was the detection success ratio. In this article, 12 of 13cases were successfully detected and the failed cases were unable to determine the fetal genotype due to insufficient number of informative SNPs. In our research, we used the coding region and highly heterozygous SNPs with 1M flanking sequence of *MMACHC* gene to construct the haplotype, and the number of informative SNPs were sufficient to build the fetal haplotype based on HMM model. Another limitation was that the phasing block size using 10x Genomics linked reads. The block size was affected by the integrity of input DNA and rigorous experimental operation. The phasing block stridden across the target gene and sufficient SNPs surrounding this region coexisted to ensure the success rate. In summary, 10x Genomics linked the reads have solved the problem of not relying on the precursor, but the success rate, high cost and complex operation restricted its clinical application. Further study is needed to develop cost-effective and simplified technology for NIPT of monogenic disease.

In conclusion, we demonstrated the feasibility of haplotype-based NIPT of MMA by sequencing a much smaller region. The sequencing depth of plasma and the number of informative SNPs were 200x and 20x, respectively. The proposition couples who have been diagnosed as monogenic carriers and have an affected child, our method is applicable to assess the repregnant fetal genotype to clinical intervention. We here provided the evidence that the same approach can be applied to other autosomal-recessive disorders, and the overall sensitivity and specificity was 100% on 78 patients tested from the previous and our study.

## Acknowledgments

We thank the patient families and physicians for cooperation during this study. This study was supported by Major Technical Innovation Project of Hubei Province (2017ACA097).

## Author Contributions

L.H., J.Y., W.Q., H.Z., L.L., and F.G, obtained patient materials; Y.W., H.W., and X.J. conducted gene analysis in blood or amniotic fluid cell DNA; C.C., J.S. and Z.P. designed the study and drafted the paper. Y.W., F.G. and W.L analyzed the data, interpreted the data and wrote a part of the paper.

## Data Availability

The authors affirm that all data necessary for confirming the conclusions of the article are present within the article, figures, and tables. Supplemental materials reported in this study are also available in the CNGB Nucleotide Sequence Archive (CNSA, ftp://ftp.cngb.org/pub/CNSA/CNP0000164/Supplementation/).

## References

1. Carrillo-Carrasco, N., R. J. Chandler and C. P. Venditti, 2012 Combined methylmalonic acidemia and homocystinuria, cblC type. I. Clinical presentations, diagnosis and management. J Inherit Metab Dis 35: 91–102.

2. Wang, F., L. Han, Y. Yang, X. Gu, J. Ye et al., 2010 Clinical, biochemical, and molecular analysis of combined methylmalonic acidemia and hyperhomocysteinemia (cblC type) in China. J Inherit Metab Dis 33 Suppl 3: S435–442.

3. Han, B., Z. Cao, L. Tian, H. Zou, L. Yang et al., 2016 Clinical presentation, gene analysis and outcomes in young patients with early-treated combined methylmalonic acidemia and homocysteinemia (cblC type) in Shandong province, China. Brain Dev 38: 491–497.

4. Evans, M. I., E. L. Krivchenia, R. J. Wapner and R. Depp, 3rd, 2002 Principles of screening. Clin Obstet Gynecol 45: 657–660; discussion 730-652.

5. Mujezinovic, F., and Z. Alfirevic, 2007 Procedure-related complications of amniocentesis and chorionic villous sampling: a systematic review. Obstet Gynecol 110: 687–694.

6. Lo, Y. M., M. S. Tein, T. K. Lau, C. J. Haines, T. N. Leung et al., 1998 Quantitative analysis of fetal DNA in maternal plasma and serum: implications for noninvasive prenatal diagnosis. Am J Hum Genet 62: 768–775.

7. Fan, H. C., Y. J. Blumenfeld, U. Chitkara, L. Hudgins and S. R. Quake, 2008 Noninvasive diagnosis of fetal aneuploidy by shotgun sequencing DNA from maternal blood. Proc Natl Acad Sci U S A 105: 16266–16271.

8. Xu, Y., X. Li, H. J. Ge, B. Xiao, Y. Y. Zhang et al., 2015 Haplotype-based approach for noninvasive prenatal tests of Duchenne muscular dystrophy using cell-free fetal DNA in maternal plasma. Genet Med 17: 889–896.

9. Meng, M., X. Li, H. Ge, F. Chen, M. Han et al., 2014 Noninvasive prenatal testing for autosomal recessive conditions by maternal plasma sequencing in a case of congenital deafness. Genet Med 16: 972–976.

10. Lam, K. W., P. Jiang, G. J. Liao, K. C. Chan, T. Y. Leung et al., 2012 Noninvasive prenatal diagnosis of monogenic diseases by targeted massively parallel sequencing of maternal plasma: application to beta-thalassemia. Clin Chem 58: 1467–1475.

11. Ma, D., H. Ge, X. Li, T. Jiang, F. Chen et al., 2014 Haplotype-based approach for noninvasive prenatal diagnosis of congenital adrenal hyperplasia by maternal plasma DNA sequencing. Gene 544: 252–258.

12. Ye, J., C. Chen, Y. Yuan, L. Han, Y. Wang et al., 2018 Haplotype-based Noninvasive Prenatal Diagnosis of Hyperphenylalaninemia through Targeted Sequencing of Maternal Plasma. Sci Rep 8: 161.

13. You, Y., Y. Sun, X. Li, Y. Li, X. Wei et al., 2014 Integration of targeted sequencing and NIPT into clinical practice in a Chinese family with maple syrup urine disease. Genet Med 16: 594–600.

14. Chen, M., S. Lu, Z. F. Lai, C. Chen, K. Luo et al., 2017 Targeted sequencing of maternal plasma for haplotype-based non-invasive prenatal testing of spinal muscular atrophy. Ultrasound Obstet Gynecol 49: 799–802.

15. Ma, D., Y. Yuan, C. Luo, Y. Wang, T. Jiang et al., 2017 Noninvasive prenatal diagnosis of 21-Hydroxylase deficiency using target capture sequencing of maternal plasma DNA. Sci Rep 7: 7427.

16. Benn, P., H. Cuckle and E. Pergament, 2012 Genome-wide fetal aneuploidy detection by maternal plasma DNA sequencing. Obstet Gynecol 119: 1270; author reply 1270-1271.

17. Norton, M. E., H. Brar, J. Weiss, A. Karimi, L. C. Laurent et al., 2012 Non-Invasive Chromosomal Evaluation (NICE) Study: results of a multicenter prospective cohort study for detection of fetal trisomy 21 and trisomy 18. Am J Obstet Gynecol 207: 137 e131–138.

18. Dar, P., K. J. Curnow, S. J. Gross, M. P. Hall, M. Stosic et al., 2014 Clinical experience and follow-up with large scale single-nucleotide polymorphism-based noninvasive prenatal aneuploidy testing. Am J Obstet Gynecol 211: 527 e521–527 e517.

19. Lun, F. M., N. B. Tsui, K. C. Chan, T. Y. Leung, T. K. Lau et al., 2008 Noninvasive prenatal diagnosis of monogenic diseases by digital size selection and relative mutation dosage on DNA in maternal plasma. Proc Natl Acad Sci U S A 105: 19920–19925.

20. Guissart, C., C. Dubucs, C. Raynal, A. Girardet, F. Tran Mau Them et al., 2017 Non-invasive prenatal diagnosis (NIPD) of cystic fibrosis: an optimized protocol using MEMO fluorescent PCR to detect the p.Phe508del mutation. J Cyst Fibros 16: 198–206.

21. Chitty, L. S., S. Mason, A. N. Barrett, F. McKay, N. Lench et al., 2015 Non-invasive prenatal diagnosis of achondroplasia and thanatophoric dysplasia: next-generation sequencing allows for a safer, more accurate, and comprehensive approach. Prenat Diagn 35: 656–662.

22. Chen, Y., Y. Liu, B. Wang, J. Mao, T. Wang et al., 2016 Development and validation of a fetal genotyping assay with potential for noninvasive prenatal diagnosis of hereditary hearing loss. Prenat Diagn 36: 1233–1241.

23. Han, M., Z. Li, W. Wang, S. Huang, Y. Lu et al., 2017 A quantitative cSMART assay for noninvasive prenatal screening of autosomal recessive nonsyndromic hearing loss caused by GJB2 and SLC26A4 mutations. Genet Med 19: 1309–1316.

24. Hui, W. W., P. Jiang, Y. K. Tong, W. S. Lee, Y. K. Cheng et al., 2017 Universal Haplotype-Based Noninvasive Prenatal Testing for Single Gene Diseases. Clin Chem 63: 513–524.

25. Huemer, M., B. Simma, B. Fowler, T. Suormala, O. A. Bodamer et al., 2005 Prenatal and postnatal treatment in cobalamin C defect. J Pediatr 147: 469–472.

26. Trefz, F. K., D. Scheible, G. Frauendienst-egger, M. Huemer, T. Suomala et al., 2016 Successful intrauterine treatment of a patient with cobalamin C defect. Mol Genet Metab Rep 6: 55–59.

